# Combining Magnetoencephalography with Telemetric Streaming of Intracranial Recordings and Deep Brain Stimulation – a Feasibility Study

**DOI:** 10.1101/2023.07.17.549294

**Authors:** Mansoureh Fahimi Hnazaee, Matthias Sure, George O’Neill, Gaetano Leogrande, Alfons Schnitzler, Esther Florin, Vladimir Litvak

## Abstract

The combination of subcortical Local Field Potential (LFP) recordings and stimulation with Magnetoencephalography (MEG) in Deep Brain Stimulation (DBS) patients enables the investigation of cortico-subcortical communication patterns and provides insights into DBS mechanisms. Until now, these recordings have been carried out in post-surgical patients with externalised leads. However, a new generation of telemetric stimulators makes it possible to record and stream LFP data in chronically implanted patients. Nevertheless, whether such streaming can be combined with MEG has not been tested.

In the present study, we tested the most commonly implanted telemetric stimulator – Medtronic Percept PC with a phantom in three different MEG systems: two cryogenic scanners (CTF and MEGIN) and an experimental Optically Pumped Magnetometry (OPM)-based system.

We found that when used in combination with the new SenSight segmented leads, Percept PC telemetric streaming only generates band-limited interference in the MEG at 123Hz and harmonics. However, the ‘legacy streaming mode’ used with older lead models generates multiple, dense artefact peaks in the physiological range of interest (below 50Hz).

The effect of stimulation on MEG critically depends on whether it is done in bipolar (between two contacts on the lead) or monopolar (between a lead contact and the stimulator case) mode. Monopolar DBS creates severe interference in the MEG as previously reported. However, we found that the OPM system is more resilient to this interference and could provide artefact-free measurements at least for limited frequency ranges.

A resting measurement in the MEGIN system from a Parkinson’s patient implanted with Percept PC and subthalamic SenSight leads revealed artefact patterns consistent with our phantom recordings. Moreover, analysis of LFP-MEG coherence in this patient showed oscillatory coherent networks consistent in their frequency and topography with those described in published group studies done with externalised leads.

In conclusion, Percept PC telemetric streaming with SenSight leads is compatible with MEG. Furthermore, OPM sensors could provide additional new opportunities for studying DBS effects.

## 1. Introduction

The network mechanisms underlying deep brain stimulation (DBS) effects for neurological and psychiatric disorders are largely unknown. One line of research aiming to elucidate these mechanisms is the characterisation of cortico-subcortical interactions and their modulation by behaviour, pharmacological agents, and DBS. Such insights can be provided by combining subcortical local field potential (LFP) recordings from DBS electrodes with simultaneous magnetoencephalography (MEG). Previous studies have already tested the technical feasibility of such recordings (Litvak et al., 2010; Oswal et al., 2016b), as well as proposed ways of effective suppression of DBS artefacts in MEG and EEG data (Abbasi et al., 2016; Lio et al., 2018; Kandemir et al., 2020).

A series of studies combining intracranial recordings from subcortical DBS targets and MEG revealed distinct cortico-subcortical coherent networks that were differentially modulated by tasks, medication and clinically effective DBS. Some of these effects were significantly correlated with disease severity and clinical improvement. A different type of MEG studies examined the effect of DBS on brain oscillations, cortico-muscular coherence or evoked responses without LFP recordings. See (Litvak et al., 2021) for a comprehensive review of many of these studies.

In most published studies with LFP recordings, these were done via temporarily externalised leads either intraoperatively or during a short interval between lead implantation in the head, and Implantable Pulse Generator (IPG) implantation in the chest. During this time, the placement of electrodes or on-scalp MEG sensors is limited due to the open wounds and oedema induced by surgery. Additionally, there is a possible temporary post-operative amelioration of clinical symptoms a phenomenon called ‘the stun effect’ (Chen et al., 2006; Eusebio and Brown, 2009; Sure et al., 2023). Therefore, the post-operative state is not fully representative of the patient’s chronic condition.

Medtronic Percept PC is the first widely available IPG which is capable of sensing LFPs while simultaneously delivering stimulation (Cummins et al., 2021; Jimenez-Shahed, 2021; Thenaisie et al., 2021). The Percept PC is a second-generation bidirectional neural interface which allows for wireless sensing and storing of brain activity using BrainSense technology. Data streaming is enabled by the connection of the IPG to an external ‘communicator’ device. The communicator initializes the connection when placed near the IPG. Once communication is established, the communicator can be placed up to a few meters away from the IPG. The communicator is then connected to a tablet via either a wireless connection or a USB cable. The tablet can be used to configure the IPG for stimulation and sensing in several different modes. We will describe below only the modes relevant to our study and refer the reader to Medtronic technical documentation for a more comprehensive description. Note that the Percept PC IPG can be used to replace older Medtronic systems without changing the already implanted leads and extension wires which will mean that patients do not undergo additional brain surgery but instead receive the Percept PC as part of their routine clinical battery replacement procedure. In newly implanted patients, a new type of leads can be implanted which allows for directional stimulation and improved sensing capabilities. These are marketed under the brand ‘SenSight’.

Since LFP sensing is done via wireless telemetry, it is possible to stream local field potentials from chronically (long-term) implanted patients. Such an advancement overcomes the limitations of the stun effect, allows for multiple recordings of the same patient for longitudinal studies, and also enables direct assessment of stimulation effects on the characteristics of the LFP. Furthermore, as the streaming is wireless there is no longer a limitation on placing on-scalp MEG or EEG sensors and no limitation on the experimental setting and amount of movement.

However, while combining the percept PC with EEG poses no specifically uncharted challenges in terms of artefacts, this is not the case for MEG. Interferences caused by data streaming, as well as by having the neurostimulator implanted in the chest, could possibly hinder high-quality magnetoencephalographic recordings. The aim of the present study is to assess the level and spatio-temporal patterns of this interference and develop recommendations for data collection and analysis to minimise its effects on the observed physiological phenomena of interest. To this end, we designed and implemented a protocol to test a Percept PC system placed inside a MEG phantom.

Several potential sources of interference should be considered. The first one is that the IPG containing ferromagnetic components generates artefacts with the subject’s movement and breathing. These artefacts should be present even with data streaming and stimulation turned off. The second is artefacts generated by electronic components of the IPG that could be recorded in the absence of streaming, stimulation and movement. The third is artefacts stemming from the wireless communication between the IPG and the communicator during data streaming. The final source of interference is active stimulation. Stimulation can be done in two ways: bipolar – with the anode and the cathode being contacts on the lead and monopolar – with the cathode being an electrode contact and the anode IPG case in the chest. Only the monopolar mode is compatible with simultaneous LFP recording. Previous work has shown that monopolar mode is far more problematic in terms of MEG interference due to strong electrical currents flowing through the entire head and upper body rather than just in the vicinity of the electrode (Oswal et al., 2016b; Kandemir et al., 2020).

Our experimental protocol was designed to test for the contribution of each of these factors alone and in combination. In order to produce generalisable conclusions, we repeated the protocol at two sites with three different MEG systems, two traditional cryogenic systems (CTF-275 and Neuromag-MEGIN-306) and a novel wearable system based on Optically Pumped Magnetometers (OPM). Together these systems cover the commonly used sensor types: axial gradiometers (CTF), planar gradiometers (MEGIN), and magnetometers (MEGIN, OPM). OPM sensors (Tierney et al., 2019; Brookes et al., 2022) measure magnetic fields generated by the brain similarly to traditional MEG but they exploit a different physical principle and therefore have different properties in terms of noise floor, bandwidth and response to artefacts.

We hope that this work will lay the foundations for further studies on combined LFP telemetry and MEG. The collected data that we are making publicly available can be used for testing and benchmarking artefact removal techniques.

## 2. Methods

### 2.1. Experimental protocol

We will first describe the sensing and stimulating modes of the Percept PC in more details.

#### Indefinite Streaming mode

This option allows streaming of bipolar LFPs from stimulation-compatible pairs, which means contact pairs that are not immediately adjacent (such as 0-3, 1-3, and 0-2 on one hemisphere and 8-11, 9-11, and 8-10 on the other). Commonly used bipolar channel pairs (0-1, 1-2 and 2-3) can be derived by linearly combining these recordings offline. Stimulation cannot be delivered in this mode.

#### BrainSense Streaming mode

in this mode simultaneous stimulation and measurement of the LFPs using the contacts adjacent to the stimulating contact are enabled, using the BrainSense technology.

Stimulation is performed in a monopolar stimulation configuration to allow for an approximately symmetric field configuration around the stimulating contact. The stimulation artefact is therefore greatly reduced due to the common mode rejection enabled by hardware design. For the stimulation ON condition, we used settings that were on the higher end of those typically used in patients to emulate the worst-case scenario (amplitude of 5mA, pulse duration 60 us, frequency 145Hz). We stimulated on the second ring from the bottom (ring 1) and recorded from the rings above and below it (0 and 2). The stimulation frequency was higher than the most commonly used 130Hz to make the stimulation-related spectral peaks distinct from streaming-related spectral peaks (at 123Hz and harmonics, see Results). There are two ways to record in BrainSense Streaming mode with the stimulation OFF. The first way is to actually turn the stimulation off. The second is to keep it on but set the amplitude to 0 mA. These two options are not equivalent in terms of both MEG artefacts (as we will show) and recorded LFP (Hammer et al., 2022). We, therefore, tested both variants separately in our protocol.

#### Legacy vs. SenSight streaming

Our preliminary testing showed that there are two streaming protocols the Percept PC can use. These are very different in terms of the artefacts they generate as will be shown below. SenSight streaming is only possible when at least one of the leads is of SenSight type. The switching between the two modes is done implicitly, and the user interface does not indicate which one is currently in use. A software bug, which was fixed in Medtronic Comm Manager software version 1.0.1213 or later, made it possible to enter legacy mode with SenSight leads by initiating a connection in untethered mode and later connecting the tablet and communicator with a cable. In the recordings conducted in London, we took advantage of this bug to test both streaming modes without changing leads. The Düsseldorf team updated their software prior to the tests, and they could only test the SenSight mode.

#### Bipolar stimulation mode

In this mode, sensing and data streaming are disabled. The stimulation is done in bipolar stimulation configuration and is included in the experiment protocol to assess the artefact of the stimulation in the MEG data.

#### Monopolar stimulation mode

The stimulation is the same as in BrainSense with stimulation ON but sensing and data streaming disabled.

### 2.2. Experimental set up

To assess the effect of stimulation and telemetric streaming on all common MEG systems and sensor types, we conducted recordings following the same protocol at two sites: UCL Functional Imaging Laboratory (hereafter referred to as London) and University of Düsseldorf MEG Lab (hereafter referred to as Düsseldorf).

In London, we performed the testing on two different MEG systems. The cryogenic CTF 275-channel MEG system (CTF/VSM MedTech, Vancouver, Canada) employs axial gradiometers as its primary sensor type. Data were sampled at 19.2 kHz and we did not use online artefact suppression (recorded in raw mode).

OPM-MEG data were acquired in an MSR (Magnetic Shields Ltd) with internal dimensions of 438 cm x 338 cm x 218 cm, constructed from two inner layers of 1 mm mu-metal, a 6 mm copper layer, and then two external layers of 1.5 mm mu-metal. A total of 31 dual-axis OPMs (QuSpin Inc., QZFM second generation) were used in the study, exhibiting a sensitivity of ∼15 fT/√Hz between 1 and 100 Hz. The sensors were placed in a 3D-printed “scanner-cast” (Boto et al., 2017), used for one of the human participants, which was built based on their structural MRI (Chalk Studios, London). Custom plastic clips were employed to organize the OPM sensor ribbon cables for effective cable management.

Sensors were positioned to evenly cover the entire head, maintaining an approximately symmetrical layout across each hemisphere. This resulted in 62 OPM data channels. Additional 11 dual axis and one triaxial OPM were not mounted on the helmet and 25 channels recorded from these sensors were not analysed. Before the start of the experiment and after each time the door was opened, the MSR was degaussed to minimise the residual magnetic field.

Data were recorded using a 16-bit precision analogue-to-digital converter (National Instruments) with a sample rate of 6 kHz. In our setup, the two sensitive OPM axes were oriented both radially and tangentially to the head, which increased the dimensionality of the data and facilitated spatial interference suppression methods (Brookes et al., 2021; Tierney et al., 2021a; Seymour et al., 2022a).

In Düsseldorf, data were acquired on a cryogenic 306-channel MEGIN system (Elekta Neuromag, Helsinki, Finland). The sensor array comprised 102 sensor triplets, each with one magnetometer and two mutually orthogonal planar gradiometers. Data were sampled at 5 kHz. Offline artefact reduction methods commonly used in this system (MaxFilter/SSS, (Taulu and Simola, 2006)) were not applied (see Discussion).

In London, a CTF current dipole phantom was used (CTF/VSM MedTech, Vancouver, Canada). The phantom consisted of a saline-filled spherical vessel placed on a plastic stand. The sphere features an array of openings at the bottom, allowing for dipole placement in various positions. We utilised the sphere inverted, with the openings facing upwards, enabling easy insertion of DBS leads and other wires without sealing the openings. Although this prevented us from positioning the sphere at the top of the MEG helmet or head cast, we believe it is not critical in this case. Two leads were inserted into the phantom on opposite sides, and their other ends were connected to the IPG via extension wires, as would be done in a patient. To close the loop for stimulation, we employed an EEG electrode fixed to the IPG case using Blu Tack adhesive. The electrode was connected to another electrode inserted into the phantom, which was submerged in saline.

For the CTF recording, the phantom was positioned inside the MEG helmet as deep as possible. For the OPM recording, the scanner cast was placed on top of the phantom. The IPG was kept at a distance roughly equivalent to the distance between the chest and head in an adult (∼33cm). For the blocks involving movement, an experimenter, present inside the shielded room throughout the experiment picked up the IPG and moved it approximately 1-2 cm back and forth, in sync with their own breathing, to simulate breathing movement.

The communicator was positioned as far from the MEG sensors as possible: ∼2m away in the CTF room and ∼3m away in the OPM room. It was connected to a tablet outside the shielded room using a cable. There was no need to move the communicator closer to the IPG to establish communication, as the shielded rooms provided a very low-noise environment.

Recordings in Düsseldorf were done using a phantom in the form of a human skull, which was filled with saline solution. The DBS electrodes, as well as other cables, could be inserted via holes in the phantom surface. The phantom was placed on a box, filled with air rhythmically controlled via an Arduino circuit board (https://www.arduino.cc/) to simulate breathing. The Arduino and the valve were located outside the shielded room with only the hose led through the shielded room wall to the phantom.

For blocks with stimulation and no data streaming, the communication session was closed prior to the recording to prevent communication-related artefacts. In the Düsseldorf session, however, this was not always done as well as for some of the London OPM recordings.

For recordings in legacy mode, the communication session began with the tablet inside the shielded room wirelessly connected to the communicator. The tablet was then taken outside the room and connected to the cable without re-initiating the session. This resulted in software warnings, but then the session continued in legacy mode. See Figure 1 for a for a schematic overview of the experimental set up and Figure 2 for photos of the setups in the three MEG systems.

**Figure 1.**
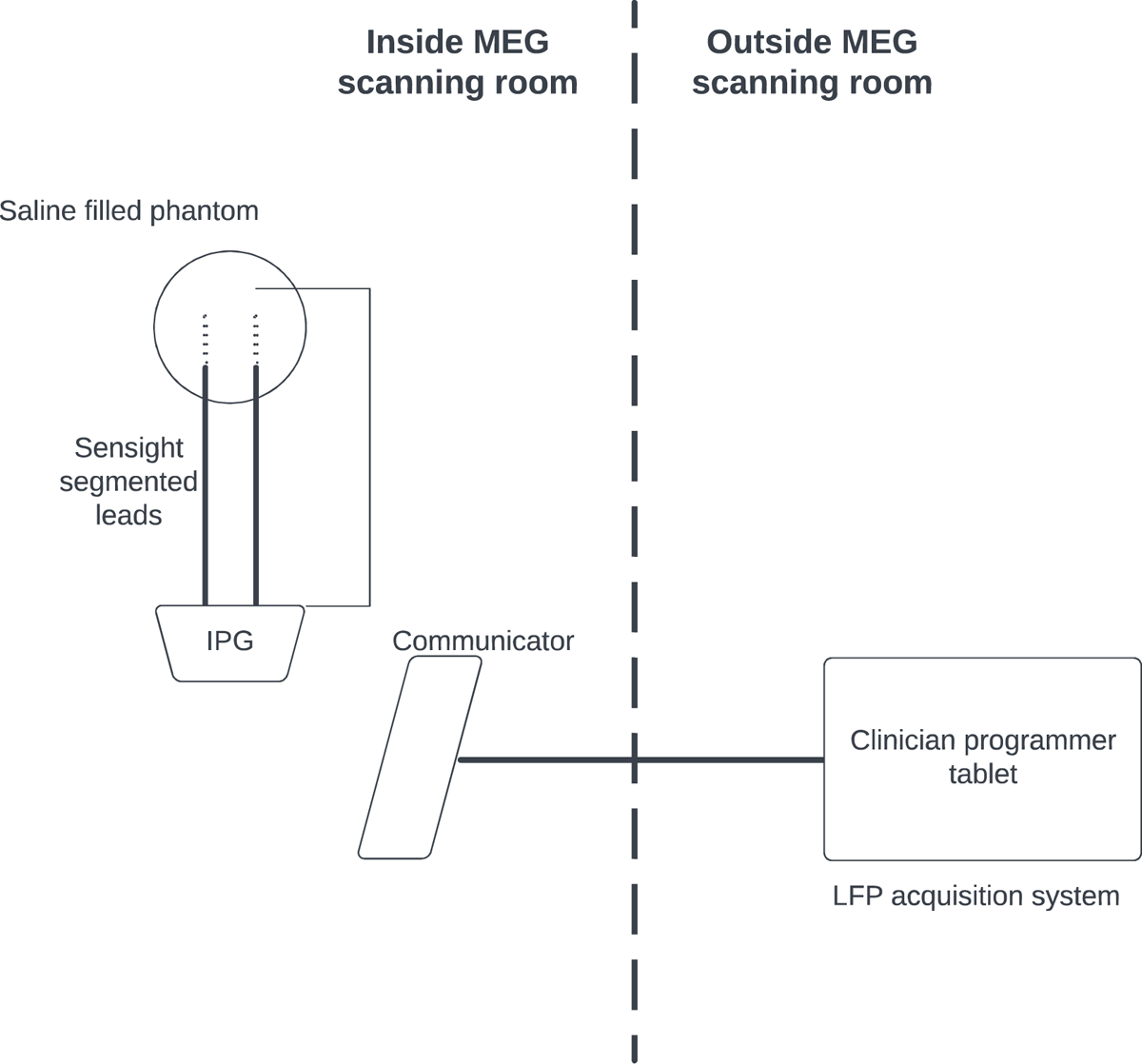
Schematic overview of our setup. Testing of the Percept PC system within the MEG scanner (either the cryogenic MEG or OP-MEG) was done using a phantom head. The phantom head and connected IPG are positioned under the MEG Dewar in the case of cryogenic MEG and under the scanner cast helmet for the OP-MEG. To minimize artefacts, the communicator is placed at a distance from the IPG once connected. The clinician programmer tablet is situated outside the shielded room and connected to the communicator via a cable fed through one of the cable guide tubes in the room walls. One experimenter can adjust various stimulation and streaming modes from outside the room, while another experimenter remains inside the room, moving the IPG during the motion conditions (London) or supervising the movement done by a pneumatic mechanism (Düsseldorf).

**Figure 2.**
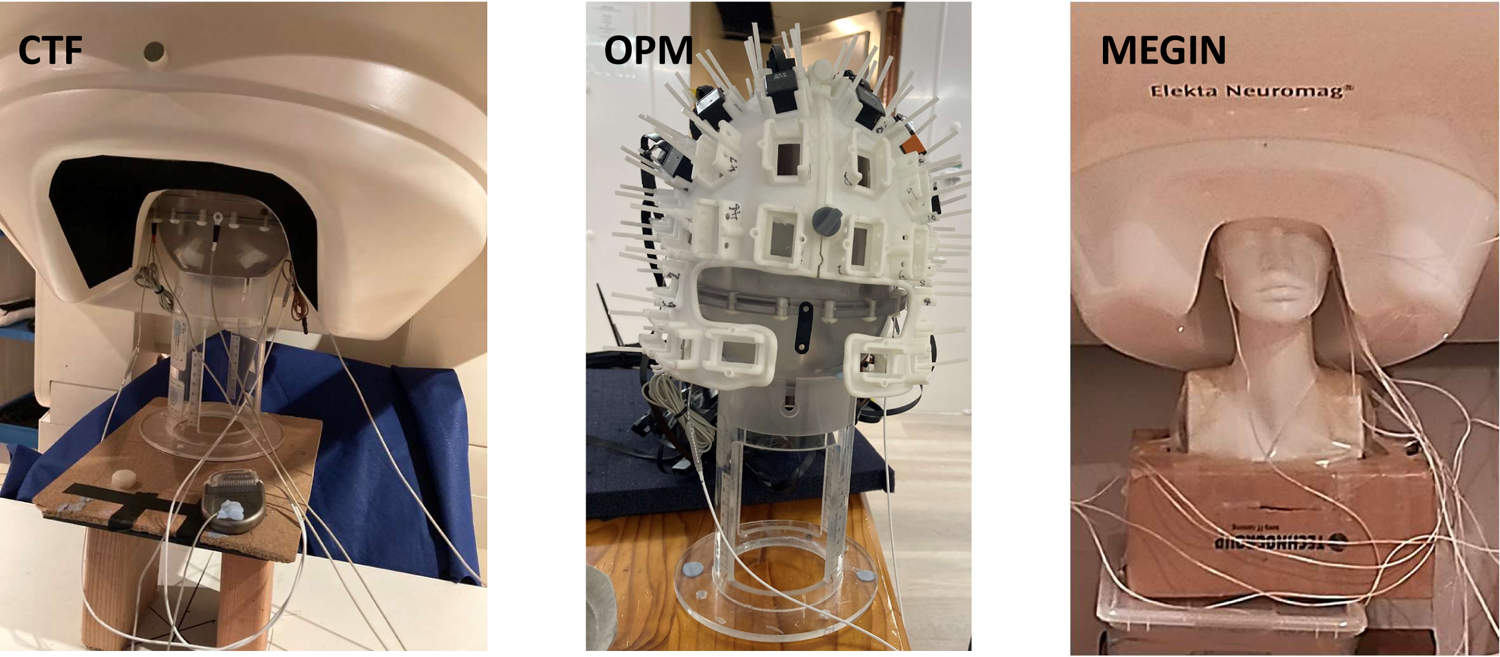
Photos of the phantom setup in the CTF, OPM, and MEGIN systems. Two SenSight segmented leads are inserted into the saline solution in the phantom and connected to the IPG via extension wires. An additional wire is also affixed to the IPG using adhesive paste, with the other end inserted into the saline solution. This arrangement simulates an electric circuit (loop) similar to what would be present in an actual patient.

**Figure 3.**
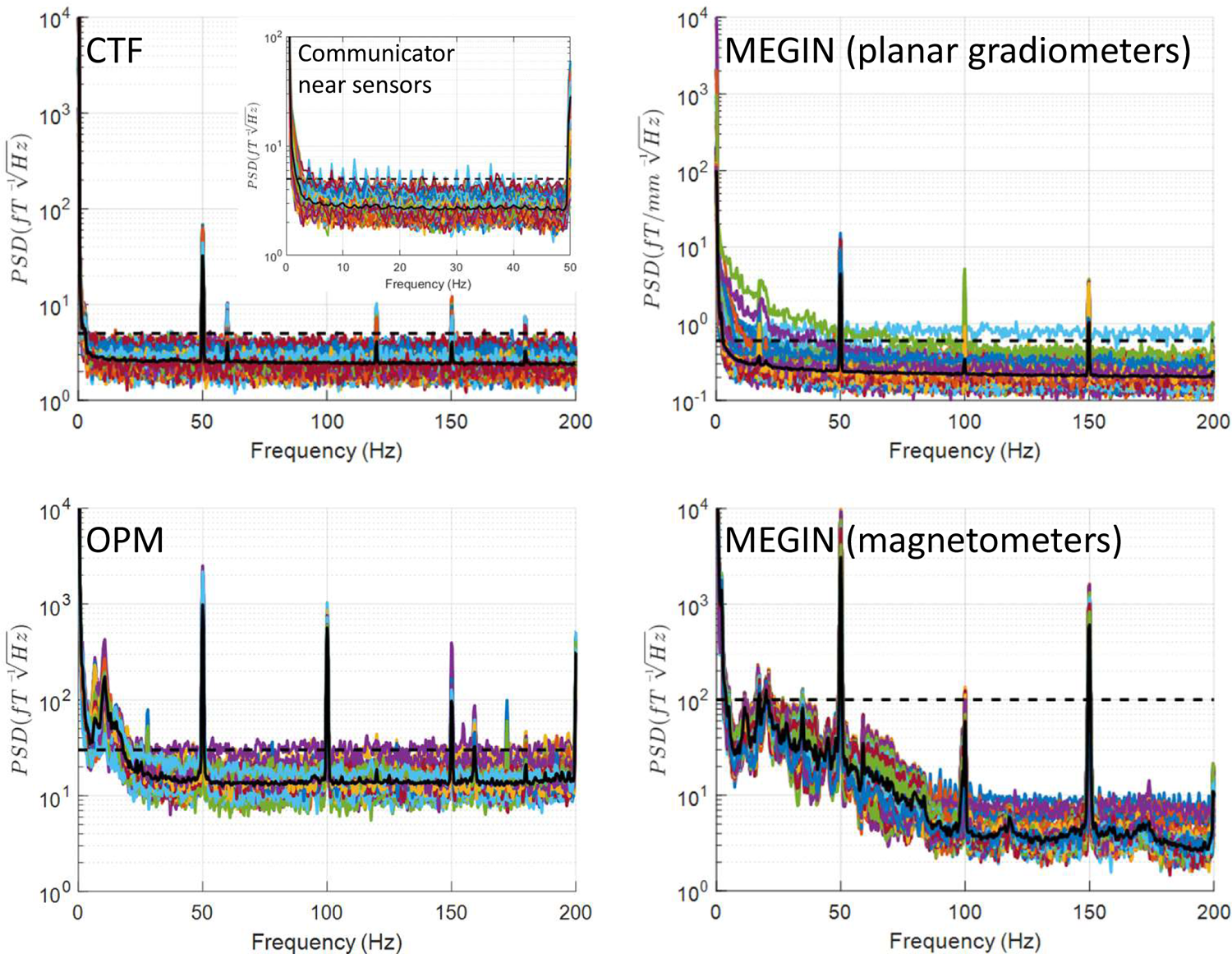
Power spectral density (PSD) for quiescent mode. The experimenter was present in the shielded room, and the IPG was connected to the phantom, but no communication session was open, and stimulation was off. The MEG spectra appear highly similar to the empty room condition (see Supplementary Figure S1), except for planar gradiometers in the MEGIN system, which exhibit an additional peak around 18 Hz and increased low-frequency noise in two sensors. In the CTF system, recordings were also conducted in this condition, with the communicator positioned closer to the MEG array (approximately 1m from the phantom). Under these circumstances, small artefact peaks were observable at 2 Hz harmonics between 4 and 48 Hz, as shown in the inset on the CTF panel.

### 2.3. Patient experiment

To confirm that our phantom results apply to real patient, we recorded a DBS patient with SenSight leads implanted in the subthalamic nucleus (STN). The data were recorded in Düsseldorf MEGIN scanner in BrainSense mode off stimulation.

### 2.4. Data analysis (phantom)

The data were analysed using Statistical Parametric Mapping (SPM, https://www.fil.ion.ucl.ac.uk/spm/). For OPM and CTF data, we only converted the data to SPM format and generated power spectral density (PSD) plots using the spm_opm_psd function with a trial length parameter of 3 seconds. MEGIN data required an additional cleaning step, which involved removing data containing flat segments and discontinuous jumps. The thresholds used for artifact detection can be found in the plot_megin.m script within the shared code.

### 2.5. Data analysis (patient)

Noise PSD for the patient recording was computed the same way as for the phantom MEGIN recording. In addition, we performed analysis akin to the one of Litvak et al. (Litvak et al., 2011) to compare coherence patterns with those previously observed in externalised recordings. The aligned MEG-LFP data were epoched to 1s segments. We then computed sensor-level coherence between the right STN channel and the magnetometer channels and tested for significance in scalp x frequency space by a parametric two-sample t-test with equal variance assumption against 10 images with the STN signal randomly permuted across trials. The significance threshold was p<0.05 family-wise error (FWE) corrected at the peak level and extent threshold of 100 voxels. We then also performed source analysis of coherence using Dynamic Images of Coherent Sources beamformer (Gross et al., 2001) implemented in the DAiSS toolbox included in SPM. Right STN was again used as the reference channel and the images were computed in the same bands as in the previous paper to facilitate comparison – alpha (7-13 Hz) and beta (15-35 Hz). SPM template head model fitted to the patient’s head shape was used to generate a corrected sphere (single shell) forward model (Nolte, 2003). A grid with 10 mm spacing was used as the source space. The cross-spectral density was computed for the planar gradiometers and regularised by reducing the dimensionality to 150 (which is a valid setting for data that was not MaxFiltered (Westner et al., 2022)). The output coherence images were z-scored and overlaid on the SPM single subject template image.

## 3. Results

Table 1 lists the conditions tested in the phantom study. As we tested several experimental factors and their combinations on three different MEG systems we have over 70 conditions in total that would be challenging to present in their entirety. Fortunately, we found no evidence that the factors we tested (e.g. stimulation and movement) interact. Also, the recordings in the different systems are largely in agreement. Therefore, it is sufficient to describe the effects of each factor separately focusing primarily on one system and mention differences between systems where present. We focus in our description on the 0-200Hz frequency range which contains most of the activity primarily studied in human MEG.

**Table 1.** The experimental protocol using different sensing and stimulation modes of the Percept PC. *<u>IPG off:</u>* There is no stimulation and the communication session is closed, *<u>Bipolar</u> <u>stimulation:</u>* This is not a sensing mode, and data will not be streamed. The stimulation is done in bipolar stimulation configuration*, <u>Monopolar stimulation:</u>* same as Bipolar stim but in monopolar configuration. The testing is done in a case positive configuration (the IPG is the positive pole)*, <u>Indefinite streaming:</u>* This option allows to stream data, recording LFPs on the electrodes in stim compatible pairs, which means contact pairs that are not immediately adjacent (such as 0-3, 1-3, and 0-2 on one hemisphere and 8-11, 9-11, 8-10 on the other). Stimulation is off during this measurement*, <u>BrainSense OFF:</u>* in this mode simultaneous stimulation and measurement of the LFPs using the contacts adjacent to the stimulating contact are enabled, using the BrainSense technology. The stimulation is done in monopolar stimulation configuration so that a geometrically symmetric configuration around the stimulating contact is possible. The stimulation artefact is, therefore, reduced due to the common mode rejection enabled by hardware design. Note that whilst stimulation in this mode is possible, one can also use it for LFP measurement with stimulation off*, <u>BrainSense ON:</u>* same as BrainSense OFF but this time the stimulation is turned on. In both the BrainSense and Indefinite streaming mode, the time domain data (i.e. LFP magnitude µV vs. time, sampled at 250Hz) is recorded as a JSON file on the clinical tablet. *<u>BrainSense ON 0:</u>* same as BrainSense ON but the stimulation amplitude is set to 0 mA. This mode is distinct from BrainSense OFF as there are switches that close for a duration equal to the pulse width and are open for the rest of the cycle. <u>Open telemetry:</u> the communication session is open but there is no stimulation and no recording. <u>BrainSense ON/OFF/ON</u>: stimulation was turned on and off several times in BrainSense mode to test for feasibility of its use as a synchronisation marker. * These conditions were not tested on the MEGIN system. ** This condition was only tested on the CTF system

| Index | Percept mode | Movement | Streaming mode |
| --- | --- | --- | --- |
| 1 | Empty room | N/A | N/A |
| 2 | IPG off | No | N/A |
| 3 | IPG off | Yes | N/A |
| 4 | Bipolar stimulation | No | N/A |
| 5 | Bipolar stimulation | Yes | N/A |
| 6 | Monopolar stimulation | No | N/A |
| 7 | Monopolar stimulation | Yes | N/A |
| 8 | Indefinite streaming | No | SenSight |
| 9 | Indefinite streaming | Yes | SenSight |
| 10* | Indefinite streaming | No | Legacy |
| 11* | Indefinite streaming | Yes | Legacy |
| 12 | BrainSense OFF | No | SenSight |
| 13 | BrainSense OFF | Yes | SenSight |
| 14* | BrainSense OFF | No | Legacy |
| 15* | BrainSense OFF | Yes | Legacy |
| 16 | BrainSense ON | No | SenSight |
| 17 | BrainSense ON | Yes | SenSight |
| 18* | BrainSense ON | No | Legacy |
| 19* | BrainSense ON | Yes | Legacy |
| 20 | BrainSense ON 0 | No | SenSight |
| 21 | BrainSense ON 0 | Yes | SenSight |
| 22* | BrainSense ON 0 | No | Legacy |
| 23* | BrainSense ON 0 | Yes | Legacy |
| 24 | Open telemetry | No | SenSight |
| 25* | Open telemetry | No | Legacy |
| 26 | BrainSense ON/OFF/ON | No | SenSight |
| 27** | IPG off, communicator near the sensors | No | N/A |

### 3.1. Empty room recording

An empty room recording, conducted without IPG and experimenter presence, provides a baseline for comparison with other conditions. Supplementary Figure S1 displays the spectra for these recordings. In the London CTF shielded room, a consistent noise level was evident across the entire range of interest, excluding extremely low frequencies near DC. The noise level’s upper limit was approximately 5 fT/√Hz, represented as a dashed line in subsequent CTF system figures. Narrowband spectral peaks occurred at line noise frequency (50Hz), its second harmonic (150Hz), and also at 60Hz and its harmonics (120Hz and 180Hz). These peaks were previously identified at this location.

The MEGIN system features two sensor types: planar gradiometers and magnetometers, each with distinct empty room noise profiles. Planar gradiometer noise remained below 0.6 fT/[mm √Hz] (indicated as a dashed line in subsequent plots) for all but one sensor, excluding low frequencies under 20Hz with higher noise and line noise frequency (50Hz) and its harmonics. Magnetometer noise stayed below 10 fT/√Hz for frequencies over 100Hz and rose to 100 fT/√Hz for frequencies under 50Hz (denoted as a dashed line in subsequent plots), with an additional increase below 5Hz. As with gradiometers, 50Hz peaks and harmonics were present.

For the OPM system, comprised solely of magnetometers, noise levels were below 30 fT/√Hz for frequencies above 30Hz (marked as a dashed line in subsequent plots). Lower frequencies exhibited higher noise, reaching 540 fT/√Hz around 10Hz. Peaks at 50Hz and harmonics were present, along with several smaller narrowband peaks at 26, 120, 155, 160, 173, and 180Hz.

The increased low-frequency noise in the magnetometer sensors, evident in both MEGIN and OPM, is likely due to the vibrations of the shielded room walls. Additionally, magnetometers, being more sensitive than gradiometers to distant sources, may be picking up ambient electromagnetic noise from the urban and hospital environment.

In the following sections, we will compare this baseline condition to various Percept PC operation modes, focusing only on spectral features absent in the empty room condition.

### 3.2. Quiescent Mode

In this condition, shown in Figure 4, both the phantom and IPG are present, but no communication session is open and stimulation is off. Although the IPG is not fully deactivated in this mode, the MEG spectra appear highly similar to the empty room condition, barring planar gradiometers in the MEGIN system, which exhibit an additional peak around 18Hz and increased low-frequency noise in two sensors.

**Figure 4.**
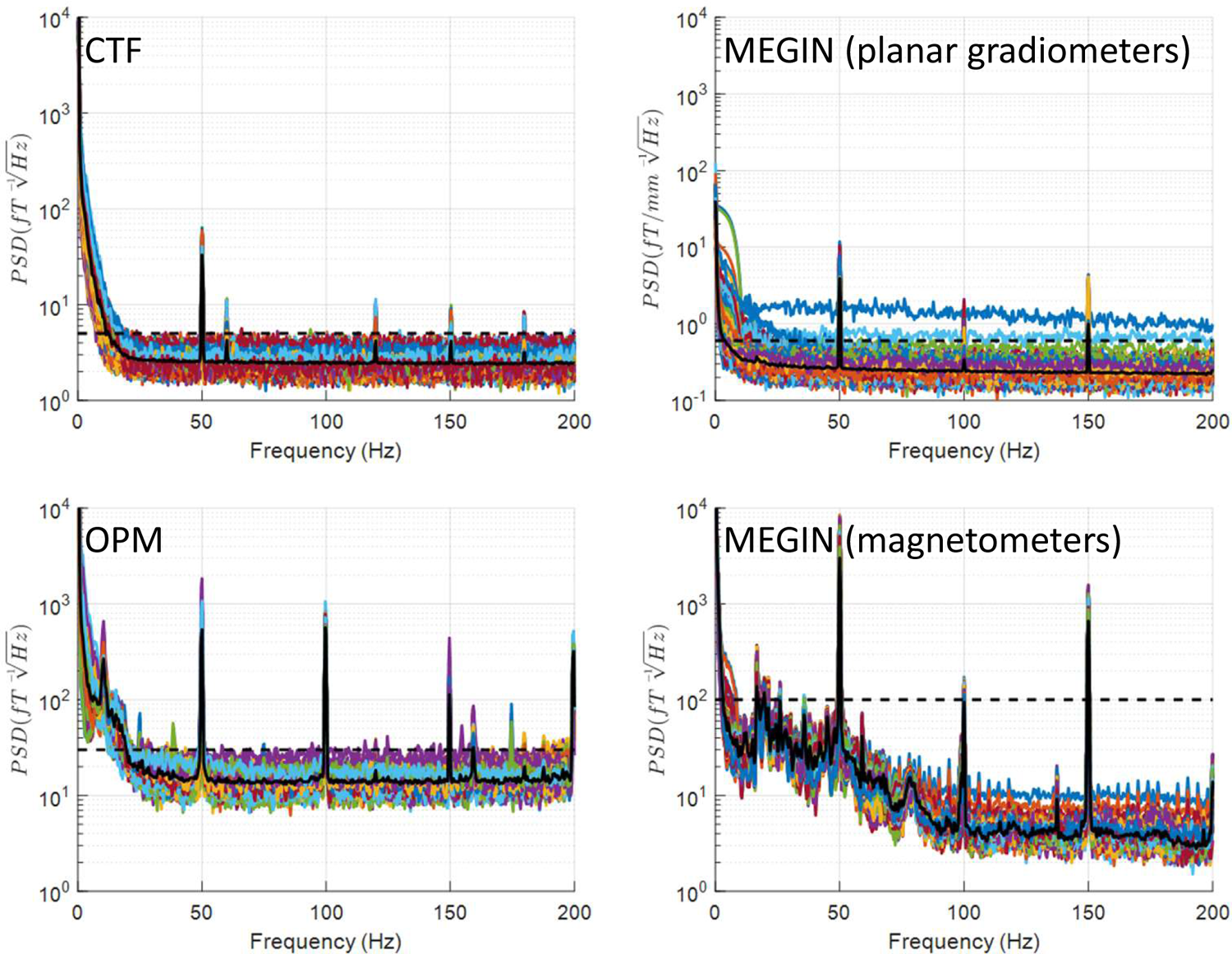
Effects of breathing-like motion of the IPG. Experimenter induced slight movement of the IPG, simulating breathing-related motion. This resulted in artefacts in low frequencies up to 20Hz. These artefacts were more discernible in the gradiometer sensors of the CTF and MEGIN systems, as these sensors exhibit lower baseline noise levels in this range. The rest of the spectrum was similar to quiescent mode recording.

In the CTF system, recordings were also conducted in this condition, with the communicator positioned closer to the MEG array (approximately 1m from the phantom). Under these circumstances, small artefact peaks were observable at 2Hz harmonics between 4 and 48 Hz.

### 3.3. Movement

Experimenter-induced slight movement of the IPG, simulating breathing-related motion, generates artefacts in low frequencies up to 20Hz, as depicted in Figure 5. These artefacts were more discernible in the gradiometer sensors of the CTF and MEGIN systems, as these sensors exhibit lower baseline noise levels in this range. To further investigate the source of these artefacts, we moved the electrodes and extension wires separately from the IPG within the MEG helmet of the CTF system while visually inspecting the signals. Wire movement did not produce any noticeable artefacts, while IPG movement caused strong deflections, confirming that the IPG is ferromagnetic but not the wires. These observations align with the weaker deflections detected when the IPG is moved further from the sensor array.

**Figure 5.**
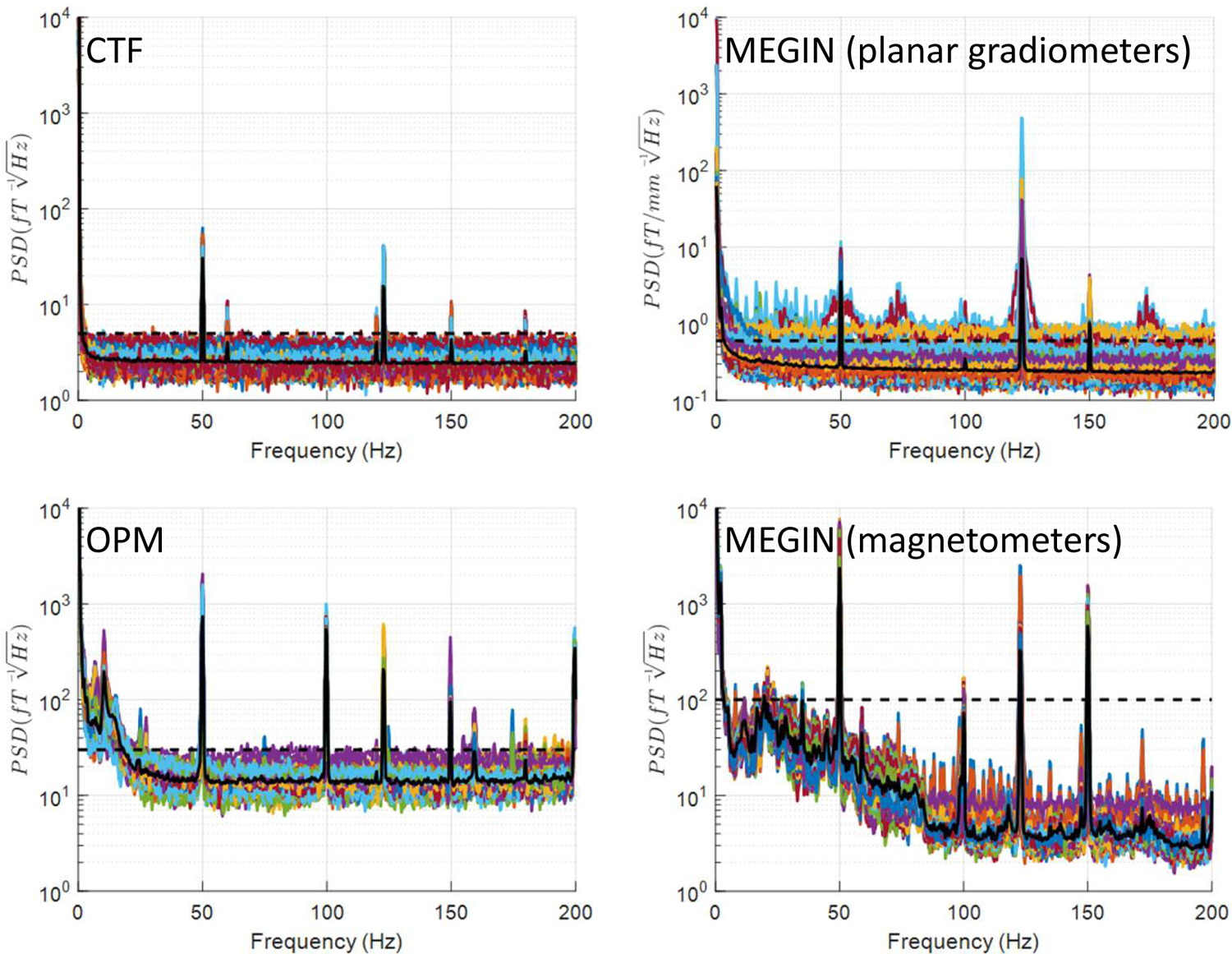
Data streaming (SenSight mode). The important and surprising finding seen here is that the noise level in this mode does not exceed that of the empty room or quiescent mode except for MEGIN gradiometers where some channels are slightly noisier. The only artefact generated by communication below 200Hz is a narrow peak at 123 Hz seen clearly on all the systems.

### 3.4. Data streaming (SenSight mode)

The primary feature emerging in this mode is a narrowband peak at 123Hz (and harmonics, which are beyond the considered range). This feature appears in all conditions involving data streaming or an open telemetry session, resulting in similar spectra for Indefinite Streaming mode, BrainSense with stimulation OFF, and open telemetry without streaming. We display the spectra for Indefinite Streaming mode in Figure 6. In some conditions, additional spectral changes occur (such as a peak at 52Hz on CTF or increased noise on some planar gradiometers in MEGIN), but these alterations are not consistently reproduced across all systems and blocks, suggesting possible unrelated environmental noise.

**Figure 6.**
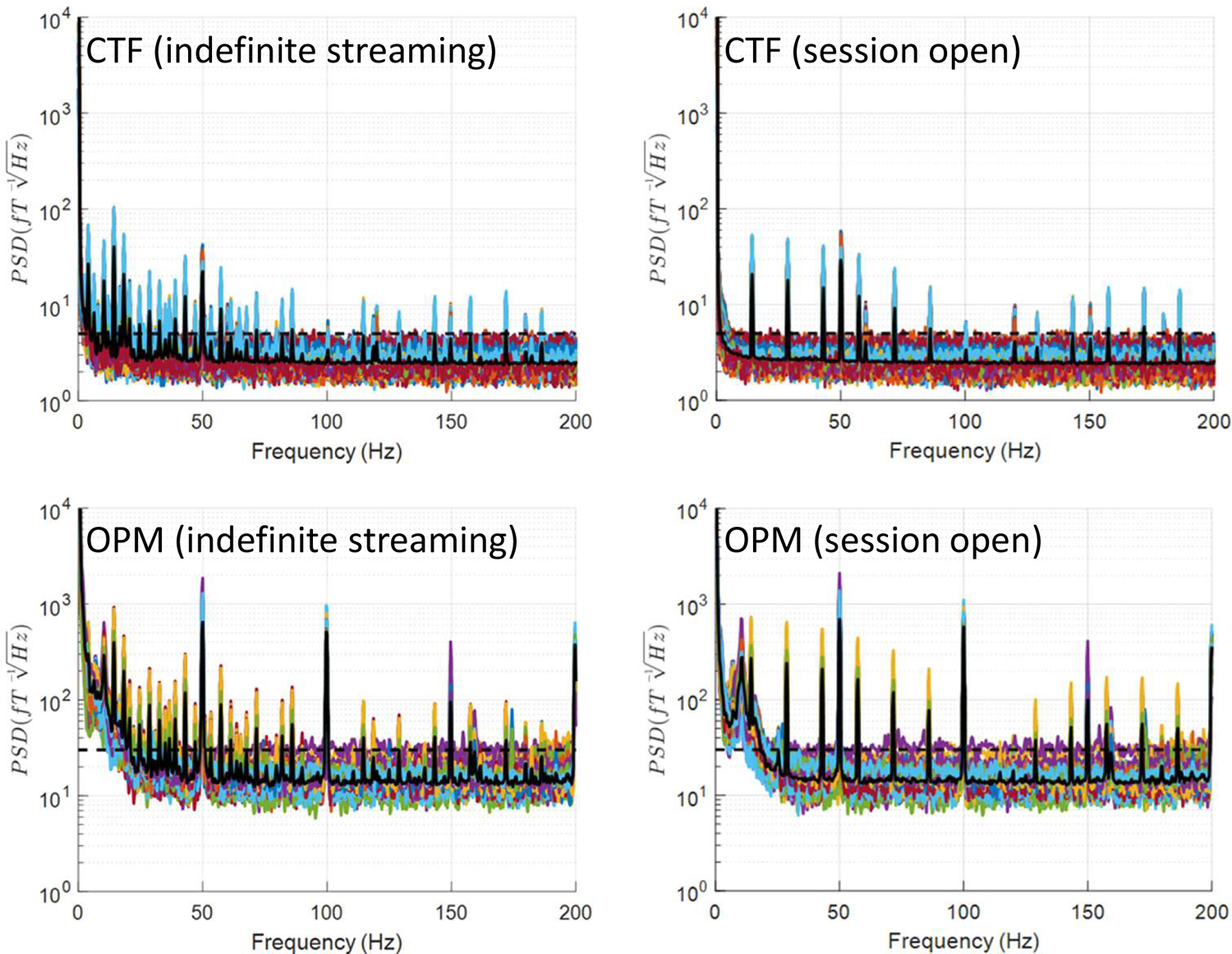
Data streaming (Legacy mode). Recordings in this mode were only done in London, therefore only results for the CTF and OPM systems are shown. During data streaming, this mode exhibits a comb-like spectrum characterized by multiple narrowband peaks covering nearly the entire range up to 150Hz and especially dense below 50Hz (left column). In the condition with open telemetry but no streaming, fewer peaks occur at 14.3Hz harmonics, with clear spectral segments between them (right column).

### 3.5. Data streaming (Legacy mode)

This mode exhibits a comb-like spectrum characterized by multiple narrowband peaks covering nearly the entire range up to 150Hz, as illustrated in Figure 6 (left column). This pattern is present for both Indefinite Streaming and BrainSense modes and appears similar in CTF and OPM data (MEGIN recordings were not conducted in Legacy mode). In the condition with open telemetry session but no streaming, fewer peaks occur at 14.3Hz harmonics, with clear spectral segments between them (Figure 7, right column).

**Figure 7.**
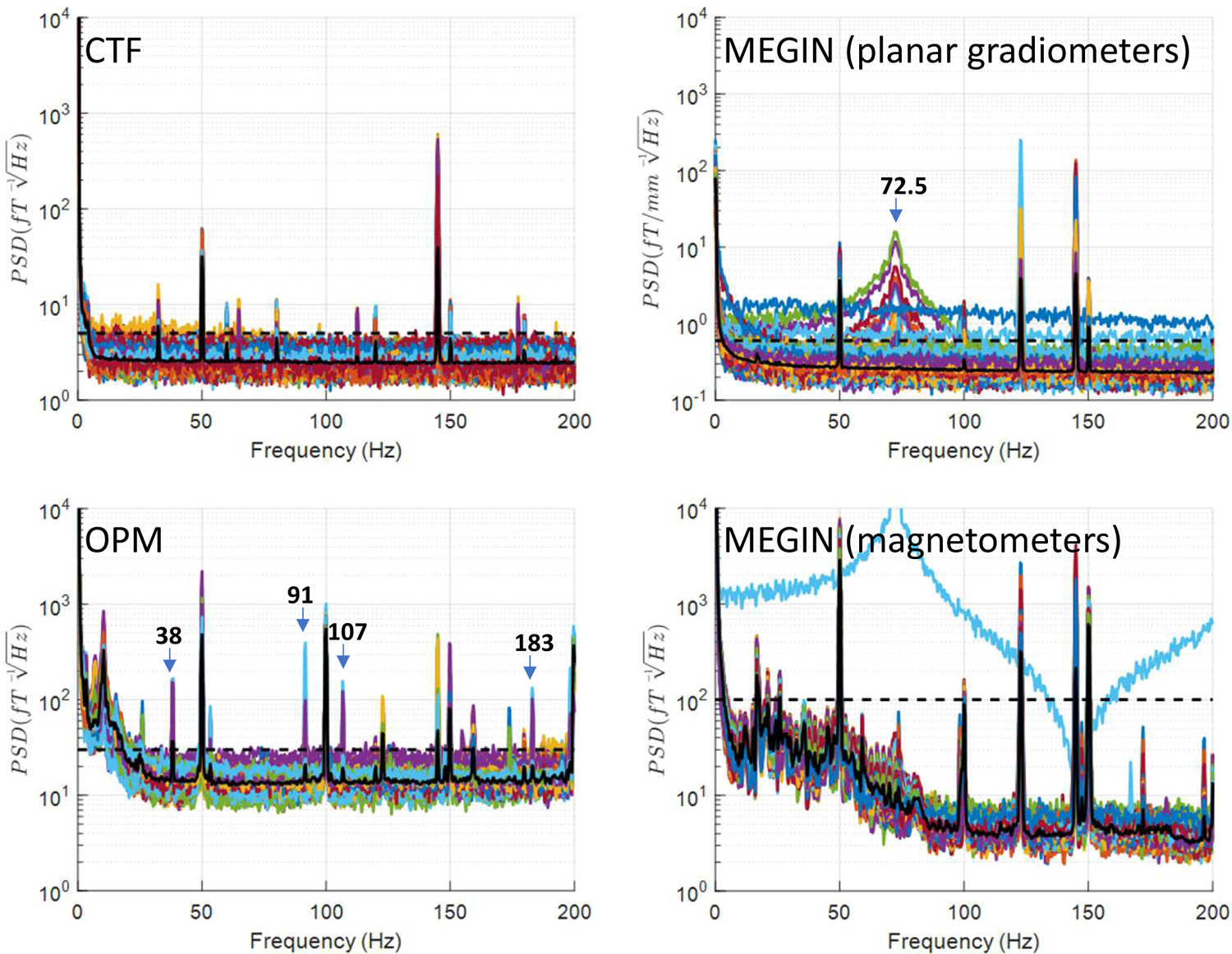
Bipolar stimulation. In this condition, a peak emerges at the stimulation frequency (145Hz) and harmonics (which are beyond the considered range). Additional spectral features vary among the three MEG systems (see Results for a detailed description). In MEGIN recordings, a 123Hz peak is also observed, likely indicating an open communication session. There is also a peak at half the stimulation frequency in some of the channels (indicated by an arrow). In OPM additional peaks appear indicated by arrows with the corresponding frequencies. See supplementary material for the discussion of the possible origins of such peaks. Crucially, for most channels and large parts of the spectrum, the noise levels do not exceed empty room levels, which is in stark contrast to the monopolar mode shown in Figure 8.

### 3.6. Bipolar stimulation

In this condition, depicted in Figure 8, a peak emerges at the stimulation frequency (145Hz) and harmonics (which are beyond the considered range). Additional spectral features vary among the three MEG systems. In the CTF system, several extra narrow-band peaks appear at 32.6Hz, 65Hz, 80Hz, 112Hz, and 177Hz. In the OPM system, extra narrow-band peaks emerge at different frequencies: 38Hz, 53Hz, 92Hz, 106Hz, and 183Hz. For MEGIN planar gradiometers, only one additional peak occurs at half the stimulation frequency (72.5Hz), but on a subset of channels, this peak is relatively wide. This peak is also present on MEGIN magnetometers, albeit less prominently. In MEGIN recordings, a 123Hz peak is also observed, likely indicating an open telemetry session.

**Figure 8.**
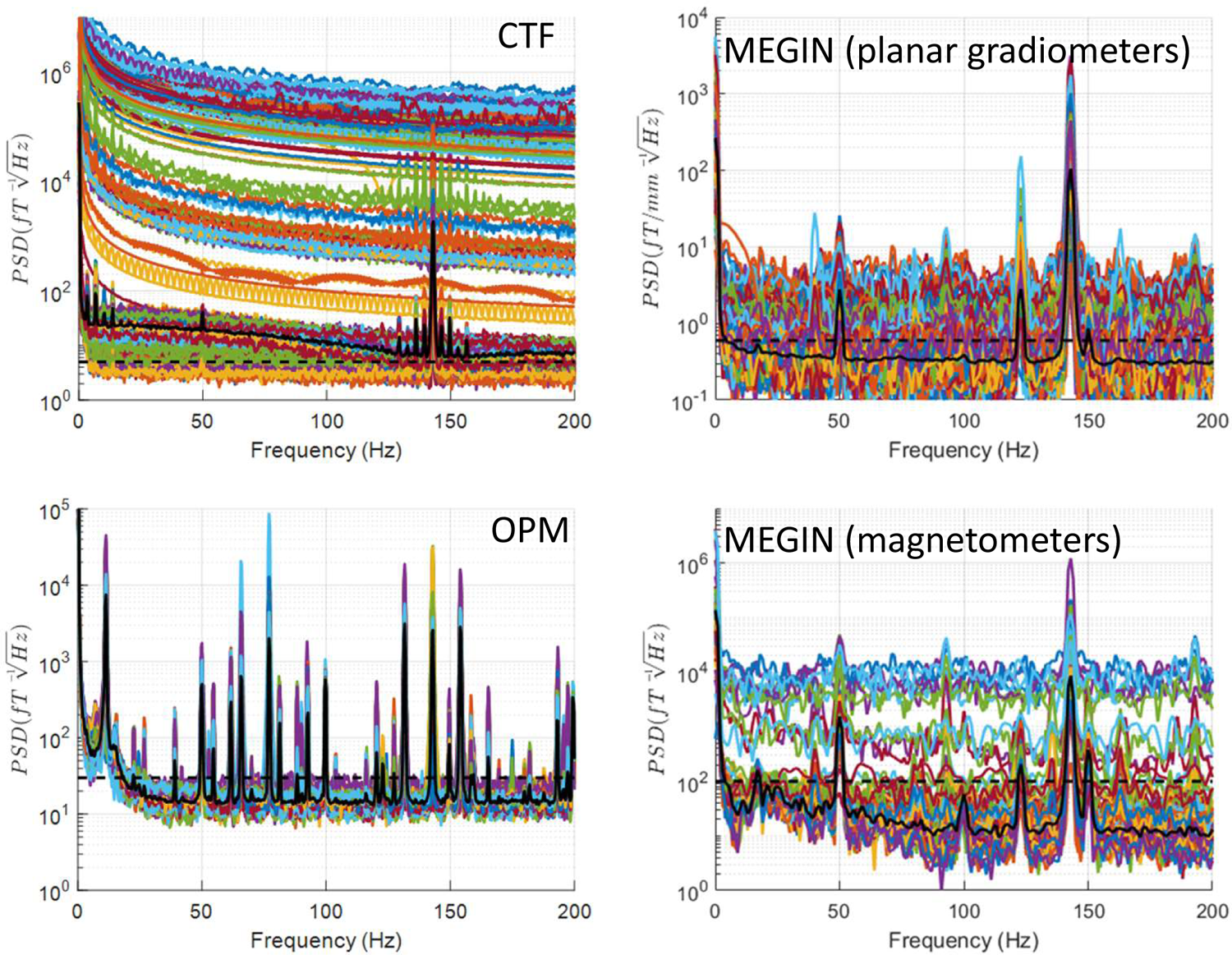
Monopolar stimulation. This mode is the most artefact-prone. In addition to the stimulation frequency peak, cryogenic MEG systems exhibit most channels exceeding the maximal empty room noise level across the entire spectrum. On the CTF system in particular, many channels display sharp jumps, which appear as wideband shifts in the frequency domain, far above the physiological signal range. Also, in the CTF data, a peak at the stimulation frequency exhibits ‘side lobes’ ranging from 16Hz below to 11Hz above the stimulation frequency. Importantly, while OPM sensors also display multiple narrowband peaks in monopolar stimulation conditions, they do not exhibit jumps, resulting in some artefact-free spectral portions that remain at the empty room noise level. See supplementary material for a more detailed analysis of the peak pattern in OPMs.

### 3.7. Monopolar stimulation

This mode shown in Figure 8 is the most artefact-prone, as previously described. In addition to the stimulation frequency, for cryogenic MEG systems, most channels exceed the maximal empty room noise level across the entire spectrum. On the CTF system in particular, many channels display sharp jumps, which appear as wideband shifts in the frequency domain, far above the physiological signal range. Also, in the CTF data, a peak at the stimulation frequency exhibits ‘side lobes’ ranging from 16Hz below to 11Hz above the stimulation frequency. This high degree of data corruption is present in all monopolar stimulation conditions, including BrainSense with stimulation on. We do not include separate figures for these conditions, but they can be generated from the data and code we share.

Importantly, while OPM sensors also display multiple narrowband peaks in monopolar stimulation conditions, they do not exhibit jumps, resulting in some artefact-free spectral portions that remain at the empty room noise level. Many of these low-frequency narrowband peaks are the result of the stimulation artefacts beating with the OPM’s modulation signal, which is set to 923 Hz (see supplementary material). Below 50Hz, OPM sensors show peaks at 39Hz, 27Hz, 25Hz, and 22Hz.

Additionally, an 11Hz peak present in empty-room data increases upon stimulation. This might have to do with the vibration at this frequency also affecting the wires between the IPG and the phantom, thereby creating a vibrating magnetic source.

### 3.8. BrainSense on Stimulation with Zero Amplitude

In this condition, streaming and monopolar stimulation are active, but the stimulation amplitude is set to zero. Consequently, in SenSight mode, there is a streaming-related peak at 123Hz and a peak at the stimulation frequency, consistent across MEG systems. Additionally, the CTF sensors display ‘side lobes’ around the stimulation frequency, similar to those observed with active stimulation, and a peak at 52Hz. Figure 9 illustrates this mode combined with movement, demonstrating that spectra for a condition combining multiple artefact sources can be accurately explained by merging the spectral features generated by each source individually.

**Figure 9.**
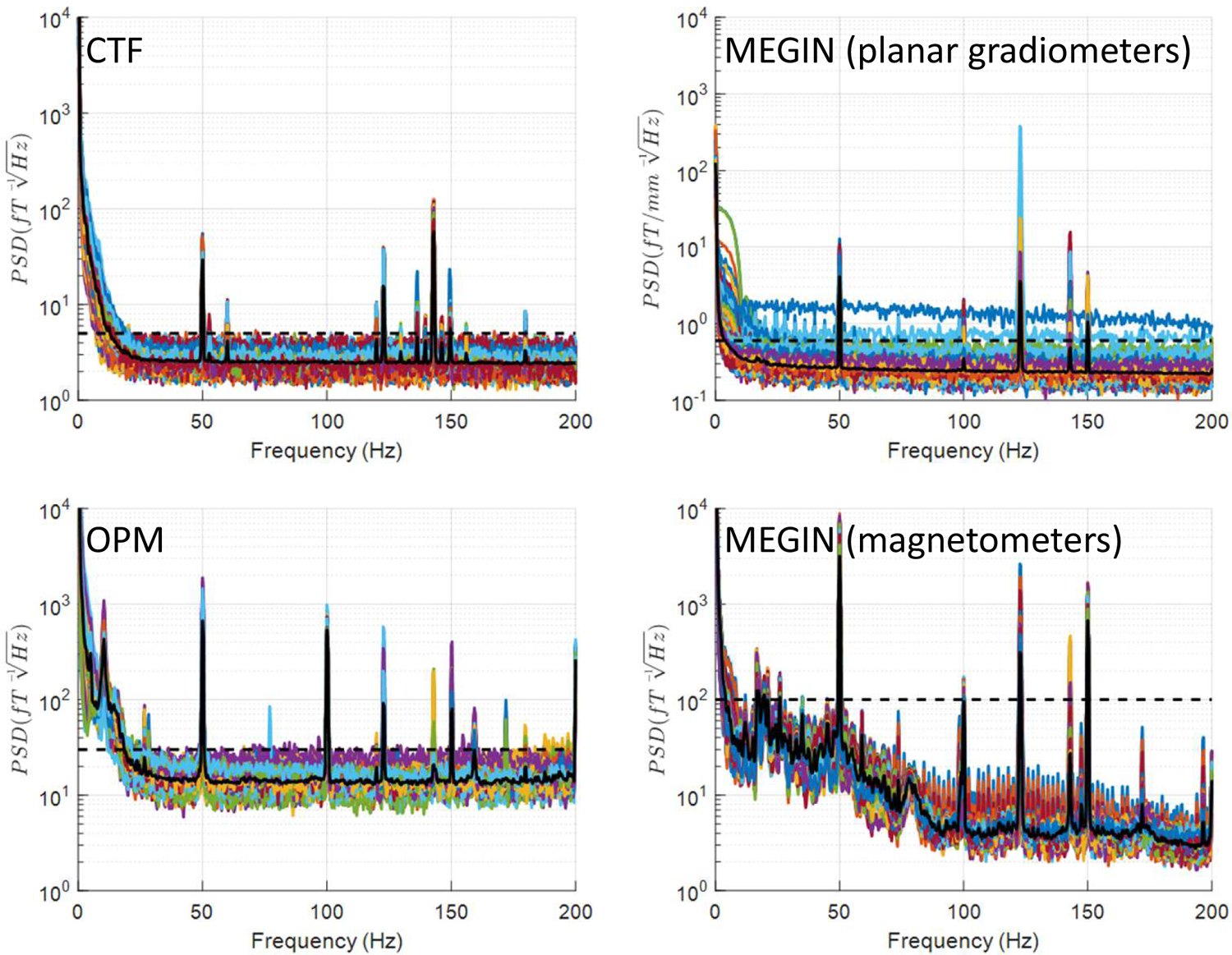
BrainSense on stimulation with zero amplitude and movement. This is an example of a ‘combined condition’ with several artefact sources which is directly comparable to patient recording shown in Figure 11. Streaming artefact at 123Hz as well as stimulation artefact at 145Hz and low frequency artefacts are present.

### 3.9. Other conditions

The other tested conditions consist of specific combinations of the factors described above, and the observed spectra can be well explained by combining the corresponding features. This supports the notion that weak or no interactions exist between these factors. We do not include separate figures for these conditions but the data and code are made available for the interested researchers to reproduce them.

### 3.10. Synchronisation

The main reason for recording subcortical LFP and MEG simultaneously is for examining measures of functional connectivity between the two signal types such as phase synchronisation. This requires precise alignment between signals recorded by the MEG system and by the IPG. As the clocks of the two systems can drift with respect to each other and the actual sampling rates can slightly differ from the specified values it is not sufficient to align based on a single marker but normally at least two markers at the beginning and the end of the recording are advised with the samples in between linearly transformed to account for the drift. The challenge specific to Percept PC and similar devices is that they do not allow adding precisely timed markers to the signals and also cannot generate triggers that can be fed into MEG. Several approaches have been suggested to circumvent this issue and we tested two of them.

#### 3.10.1. Using stimulation to synchronise

Stimulation introduces artefacts in both LFP and MEG. By switching the stimulation on and off multiple times at the start and end of a recording block, we can generate matched artefact patterns in both modalities for offline comparison. This is only feasible in BrainSense mode, which allows for stimulation. It is important to disable the ramping feature that gradually increases the stimulation amplitude over several seconds to maintain sharp boundaries at stimulation onset and offset for accurate synchronisation.

We recorded MEG signals while toggling stimulation on and off with these settings. In all three systems, the onset and offset times of stimulation are clearly discernible (Figure 10). In the CTF system, which is more severely affected by monopolar stimulation, some pre-processing was needed to clearly identify the pattern. Applying the derivative of the signal (using MATLAB’s diff function) was effective for most channels. For channels with larger jumps, taking the logarithm of the absolute value of this derivative helped bring the entire recording into a similar range.

**Figure 10.**
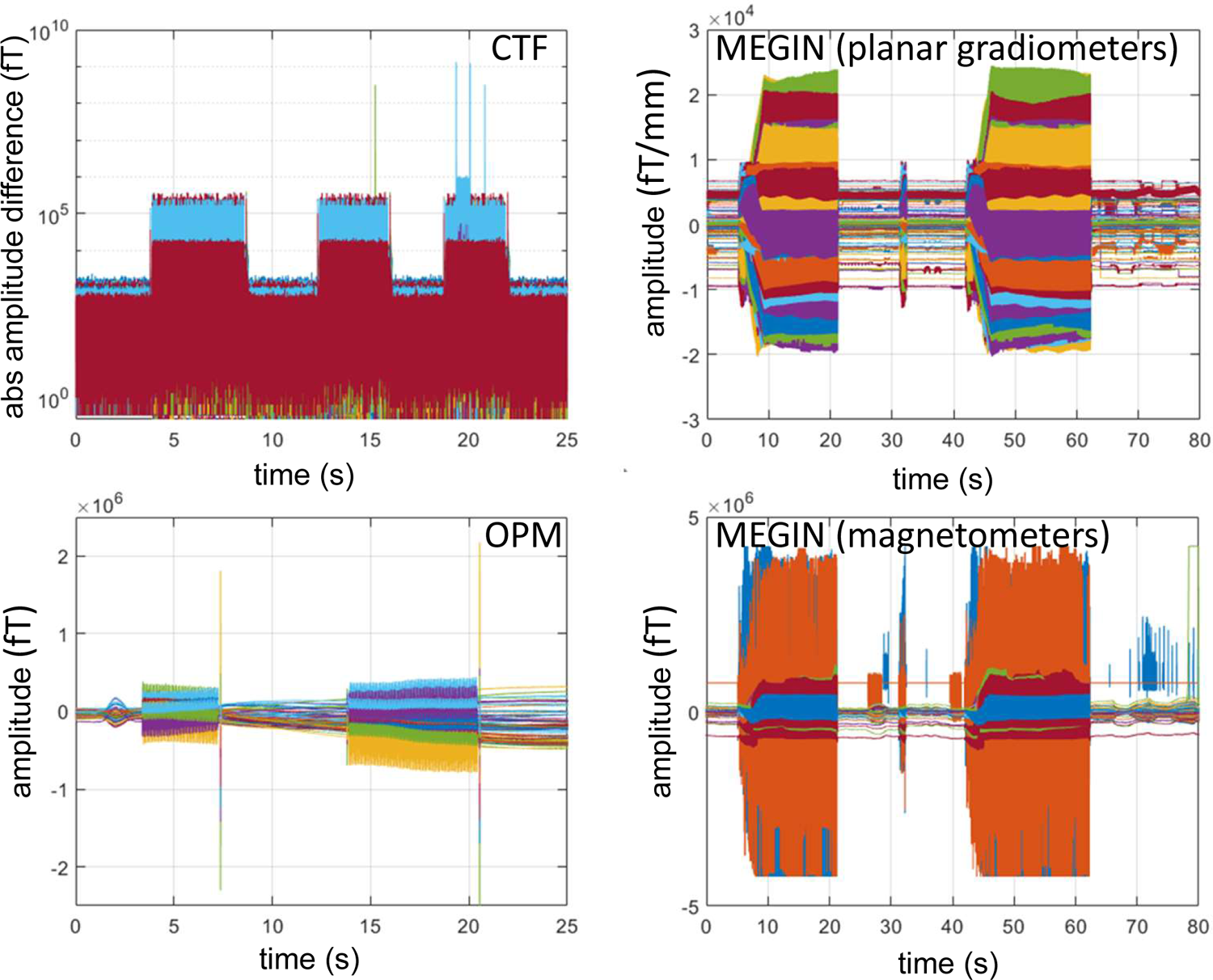
Using stimulation for synchronisation in BrainSense mode. The plots show time domain data recorded while toggling the stimulation on and off in BrainSense mode. On all the systems the onsets and offsets of stimulation are clearly visible. The CTF data required pre-processing (computing log of the absolute difference between adjacent samples) due to the presence of high-amplitude jumps. Artefacts can also be seen with the stimulation on.

#### 3.10.2. Tapping on the IPG

Another possibility to create a common signal for synchronisation is by mechanical impact on the IPG. A patient can be requested to tap with an open palm of the hand on the IPG and on an EEG electrode placed on the skin above it and this can create simultaneous artefacts in both that can be used to match the recordings offline. To assess the sensitivity of the LFP recording to mechanical impact we placed the IPG on an elastic pressure sensor, covered it with a double layer of leather and tapped on it with an open palm at varying force level, from the maximal non-painful level to a very light tap. The leads were placed in a cup with saline during this procedure and recording was performed in Indefinite Streaming mode. The recording showed clear artefacts in the LFP even for the lightest taps tested (data not shown). However, from trying this method on several patients with SenSight leads, the results were less encouraging and, in most cases, we could not identify clear artefacts. A larger number of patients needs to be examined to see if this method is viable at least in a subset of patients. But in any, case if planning to use this method one should pre-test it to see whether it works in the specific patient.

#### 3.10.3. Using cardiac artefact

In many cases Percept PC LFP recordings get contaminated by electrocardiogram (ECG) (Neumann et al., 2021; Stam et al., 2023). If the ECG signal is clear enough it can be matched to surface ECG recorded synchronised to the MEG signal as a way of syncing MEG and LFP. From our experience, standard cross-correlation on raw signals works quite well similarly to what is described for artificially generated random noise in (Oswal et al., 2016b). However, SenSight leads are less prone to ECG contamination and also new Percept PC IPG implantations are often done on the right side to further reduce ECG artefacts. Thus, this method can also potentially work only in a subset of patients and requires pre-testing.

### 3.11. Patient recording

To corroborate the conclusions drawn from our phantom testing, we conducted a recording in a Parkinson’s patient implanted with Percept PC and SenSight leads in the MEGIN system in Düsseldorf. The recording was completed in BrainSense mode without stimulation. We computed MEG spectra similarly to the phantom case, and the results (Figure 11) fully aligned with expectations based on our phantom recordings. Alongside line noise peaks, also observed in phantom and empty room recordings, we detected a communication-related peak at 123Hz and low-frequency activity below 20Hz, with a local peak at 8Hz—likely signifying physiological alpha activity.

**Figure 11.**
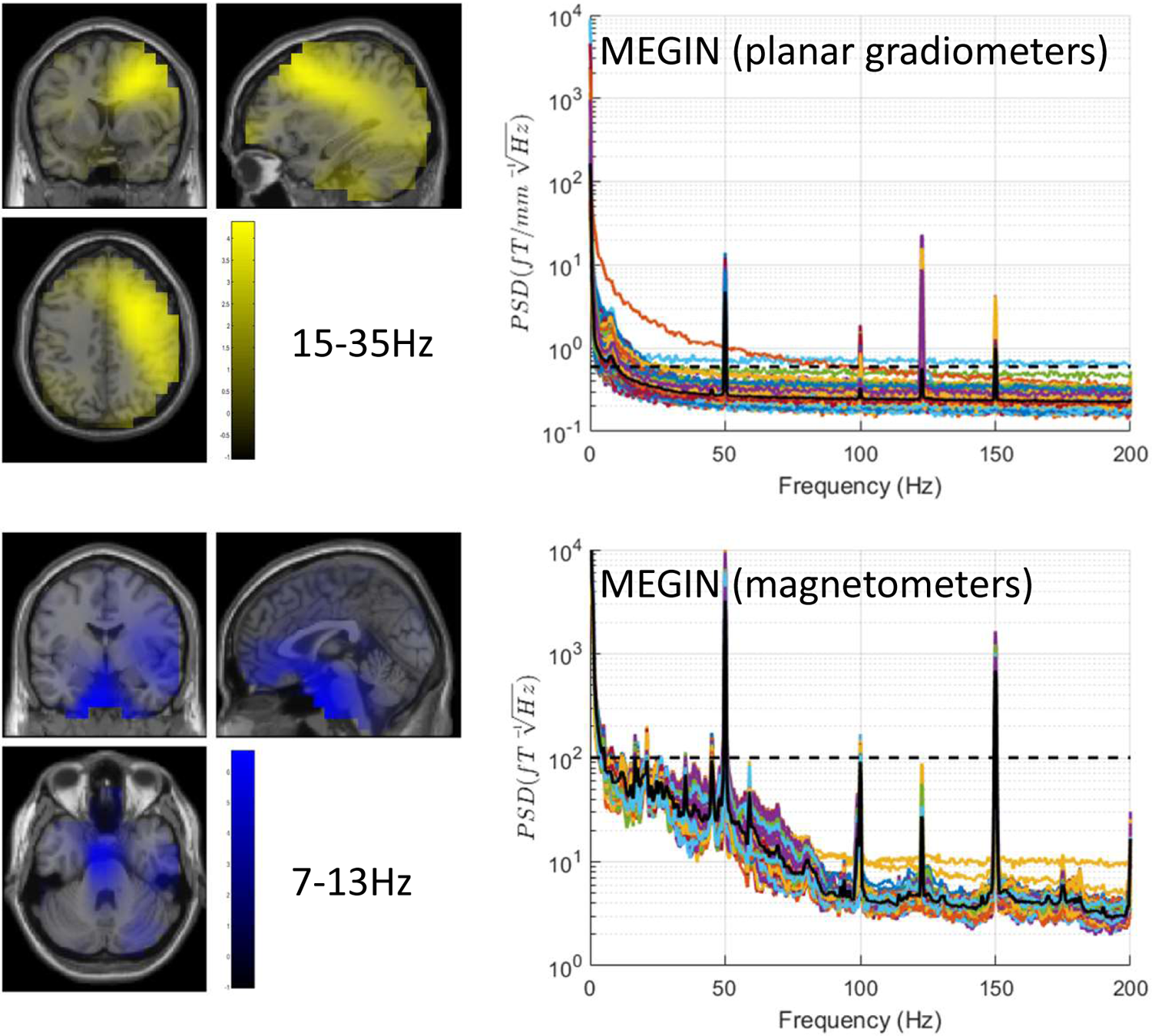
Results of patient data analysis. The recording was done in BrainSense mode off stimulation. The top left panel shows topographic image of cortico-subthalamic coherence in the beta band (15-35Hz) showing a peak in the sensorimotor areas ipsilateral to the STN. The bottom left panel is a similar localisation for the alpha band (7-13Hz) with the main peak near the brainstem and secondary peak in the ipsilateral temporal lobe. These results are remarkably consistent with previously reported group results (cf. Fig. 5 in (Litvak et al., 2011)). The right panels show power spectral densities for planar gradiometers (top) and magnetometers (bottom) with the dashed lines at the same levels as in the phantom figures. The noise levels are comparable to the corresponding phantom measurements.

We also examined significant sensor-level coherence between the right STN channel recorded on Percept PC and the MEG array, revealing three significant clusters: 4-12Hz, 6-11Hz, and 13-23Hz (not shown). Note that separate clusters could overlap in frequency due to the test being conducted on three-dimensional scalp x frequency images, allowing for overlap in frequency but separate scalp locations.

Following this, we conducted a source analysis of coherence in standard alpha (7-13Hz) and beta (15-35Hz) bands, as defined in our previous study of MEG-STN coherence in patients with externalised leads. The beta band image showed a peak in the frontal areas ipsilateral to the STN, while the alpha images displayed the largest peak near the brainstem and an additional local peak in the ipsilateral temporal lobe. These findings are in complete accordance with previous group results.

## 4. Discussion

Understanding the interference caused by the Percept PC DBS system in MEG recordings is crucial for the accurate interpretation of neural signals and the optimisation of research and clinical applications involving both technologies. As DBS has emerged as an effective therapy for several neurological and psychiatric disorders, there is an increasing need to study its underlying neural mechanisms using neuroimaging tools such as MEG. Simultaneous recordings of DBS and MEG can provide valuable insights into the real-time effects of stimulation on brain dynamics and connectivity.

Moreover, the Percept PC DBS system’s ability to record local field potentials (LFPs) directly from the implanted electrodes offers a unique opportunity to investigate the relationship between invasive and non-invasive neural signals. Combining these measurements can help advance our understanding of the pathophysiology of various disorders, as well as the mechanisms and biomarkers of DBS therapy.

However, the presence of interference in MEG recordings due to the DBS system can hinder the extraction of accurate and reliable information from these data. By characterizing and understanding these artefacts, researchers can develop effective strategies for minimizing their impact on the analysis and interpretation of MEG signals. This understanding can lead to more accurate and reliable results, ultimately improving our knowledge of brain function and the effectiveness of DBS therapy.

In our comprehensive investigation of the impact of the Percept PC DBS system on MEG recordings, we identified several consistent artefact patterns across different MEG systems and sensor types.

The main findings include:

1. Low-frequency artefacts were associated with IPG movement, reflecting the influence of breathing-related motion.
2. Wireless communication between the IPG and communicator generated distinct artefact patterns for SenSight and legacy modes. SenSight mode produced peaks at 123Hz and its harmonics, while legacy mode exhibited a comb-like pattern with multiple peaks, particularly dense below 50Hz.
3. Active DBS created artefacts at the stimulation frequency and additional frequencies that varied between MEG systems and sensor types. Bipolar stimulation generated less severe artefacts compared to monopolar stimulation, with the latter leading to severely degraded signals as previously described (Oswal et al., 2016b; Kandemir et al., 2020).
4. The communicator produced artefacts at 2Hz harmonics when positioned close to the sensor array.

Two particularly noteworthy findings for the research community are that the SenSight mode enables simultaneous streaming and LFP recording with relatively mild MEG signal contamination, and that OPMs are less severely affected by monopolar stimulation than cryogenic systems.

### 4.1. Comparison to previous studies

There are several types of previous studies that we could compare our results against. One type is simultaneous LFP and MEG recordings in patients with externalised wires (Litvak et al., 2010, 2011; Hirschmann et al., 2011). Litvak et al. (Litvak et al., 2010) described a heartbeat-locked artefact that originated from movement of ferromagnetic extension leads that were used to externalise recording contacts. The relation between the artefact and these leads was confirmed by a later phantom study reported in (Oswal et al., 2016b) and the fact that Hirschmann et al. who used custom-made non-ferromagnetic extension leads did not observe artefacts of similar magnitude and were able to record data comparable to that of subjects with no wires. Conversely, these studies did not have an IPG present so there were no low-frequency artefacts associated with breathing and movements of the torso. Based on these results, we can expect that if the wires in patients with Percept PC are non-ferromagnetic as appears to be the case from our testing, we can expect no artefacts generated inside the sensor array when the stimulation is off. Artefacts generated by movement of the IPG should be relatively easy to remove by topography-based methods as they are clearly distinct from brain signals.

The second type of studies are phantom studies that examined the effect of stimulation. Oswal et al. (Oswal et al., 2016b) focused primarily on monopolar stimulation and Kandemir et al. (Kandemir et al., 2020) – on bipolar. In line with these studies, we also got severe stimulation artefacts in the monopolar condition with relatively mild artefacts in the bipolar case. The exact patterns of subharmonic peaks we saw in our data were not observed in previous studies likely due to the different stimulators used which were an external Medtronic stimulator in the Oswal et al. study and Abbot stimulator in the Kamdemir et al. study. Also, it should be noted that in both studies the authors tried to emulate the effect of head movement so they systematically moved the phantom but not the stimulator whereas we have done the opposite to avoid further complicating the design and assuming that head movement effects will not differ from those described previously.

The third type of relevant studies are those that examined stimulation effects in patients inside the MEG such as (Airaksinen et al., 2011; Abbasi et al., 2016, 2018; Oswal et al., 2016a; Luoma et al., 2018). All the studies on this (non-exhaustive) list except the study of Oswal et al. used the MEGIN system and bipolar DBS in patients with implanted IPGs. Also, all of them employed various artefact suppression methods before analysing the data, so the results cannot be directly compared to ours. However, the one conclusion that can be drawn from these studies is that one can be optimistic about the prospects of cleaning the data sufficiently well in patients with Percept PC to see physiological effects in the MEG data as the artefacts reported by these authors were comparable to those we observed in our phantom. The study of Oswal et al. exemplifies the challenges one would face when analysing data with monopolar DBS and shows that these challenges though severe are not insurmountable. Similar challenges can be expected when using the BrainSense mode with effective DBS on Percept PC.

### 4.2. Offline artefact removal

The purpose of the present work is characterisation of the artefacts associated with the use of Percept PC in combination with different MEG systems and sensor types. To reveal the artefacts in the most comprehensive manner, we did not use any denoising methods, including ones that are routinely applied at most MEG sites, such as synthetic gradient on the CTF system (McCubbin et al., 2004), Spatiotemporal Signal Space Separation (tSSS) on MEGIN (Taulu and Simola, 2006) and similar methods recently developed and adapted for OPM systems (Seymour et al., 2022b; Tierney et al., 2022). Applying these methods and optimising them for each system and stimulation condition would require additional research that we aim to facilitate by making our phantom data freely available. Therefore, it should be noted that our results represent the worst-case scenario and it might be possible to substantially denoise the data at least for some of the cases we presented.

Previous studies have shown successful application of ICA-based methods (Abbasi et al., 2016; Kandemir et al., 2020) as well as tSSS (Airaksinen et al., 2011, 2015; Luoma et al., 2018) for suppression of artefacts of IPG movement and bipolar DBS. We are, therefore, not suggesting any particular MEG system as superior when used with Percept PC. A fair comparison would necessitate optimal configuration of the artefact suppression method specific to each system and the answer is likely to also be dependent on the paradigm and streaming mode.

### 4.3. Advantages of OPM technology over cryogenic MEG

The QuSpin OPMs used in this study operate in an open-loop mode, meaning that once the sensors are initialised, there are no active electronic systems trying to counteract the field the sensors are subjected to. This means that a sensor can go out of its prescribed dynamic range (in this case ±4.5 nT from its initialised field), but an OPM of this design can return back to its zero-point. Conversely, the superconducting quantum interference device (SQUID) sensors in the CTF and MEGIN systems used a closed-loop mode to remain in their operational range, which we believe is not able to counteract the monopolar DBS interference, leading to SQUID jumps and resets that are represented as large broadband noise in their spectra. We believe the bulk of the narrowband peaks in OPM spectra below 180 Hz are explained by beating of high-frequency components of the DBS system interacting with the modulation signal at 923 Hz (see supplementary material). For these particular OPMs, these effects are unavoidable, but for other OPMs with a different modulation frequency, it might lead to considerably fewer peaks in this part of the spectra. These findings may open up opportunities for studying DBS effects in restricted frequency bands.

### 4.4. Recommendations for future patient studies

One clear outcome of our study is that concurrent Percept PC streaming and MEG can be a viable alternative to recordings in externalised patients. This, however, comes with several caveats.

As the SenSight mode is only possible in patients with SenSight leads, patients with older leads which will comprise the majority of Percept PC patients in many centres until more new implantations are done, are not suitable for MEG-LFP studies. The problem might be finessed by an effective artefact removal method that will clean the streaming artefacts, but this is unlikely to result in high-quality MEG data, due to the large number and density of artefact peaks in the range that is of most interest (below 50Hz). Thus, combining LFP streaming in the legacy mode with MEG might not be worth the effort, especially since EEG is a viable alternative to MEG in these patients and it is not affected by streaming or ferromagnetic artefacts.

Also, Percept PC can be used for study of effects of bipolar DBS and the artefact suppression methods previously suggested for other stimulator models are likely to be effective for it as well. In this mode, however, it is not possible to record and stream LFP data.

When it comes to monopolar stimulation and BrainSense streaming on DBS, the challenges are far greater. As shown previously (Oswal et al., 2016a, 2016b) some physiologically-relevant results can be obtained from this kind of data, partially in relation to LFP-MEG coherence which is more robust to artefacts than MEG power and the same methods and conclusions are likely to apply to Percept PC studies. However, one should consider carefully whether there is justification for using MEG rather than EEG.

Using OPMs for study of monopolar DBS effects could be particularly promising, particularly when one is interested in a limited frequency range and can render this range artefact-free by adjusting the stimulation frequency and the sampling rate. OPM systems are also currently being refined for recordings during naturalistic movement and flexible sensor placement to access areas which are challenging for cryogenic MEG such as hippocampus (Tierney et al., 2021b), cerebellum (Lin et al., 2019) and spinal cord (Mardell et al., 2022). Moreover, new types of OPM sensors are still being developed which might have even better properties in terms of noise floor, bandwidth and resilience to external interference(Kowalczyk et al., 2021; Gutteling et al., 2023). Our study demonstrates the feasibility of OPM studies in Percept PC patients.

In terms of practical recommendations based on our experience, we would suggest to keep the communicator as far away from the sensor array as possible. Also, it is advised to update the tablet software to the latest version to avoid it accidentally switching to legacy mode and also visualise the data PSD after each block for quality control as the difference between legacy and SenSight modes is not easily seen by eye in the raw data. For OPMs, it is advisable to close the telemetry session when not streaming data because telemetry-related harmonics might leak into lower frequencies because of the non-linear effects present in this system. This was not, however, observed in practice in our data.

Synchronising MEG and LFP is achievable by toggling stimulation on and off in BrainSense mode. However, this method may disrupt some MEG channels for the remainder of the recording block, necessitating their exclusion from the analysis.

As for synchronisation in the Indefinite Streaming mode, a comprehensive solution is currently unavailable. Apart from the tapping and ECG methods we tested, it is possible to use a transcutaneous electrical nerve stimulation (TENS) machine to introduce simultaneous artefacts into LFP and MEG (Thenaisie et al., 2021). We have no experience with this method, and to our knowledge, it has not been tested in a MEG environment. Furthermore, even if effective, it would require ethical approval and could potentially cause additional discomfort to the patient.

The Indefinite Streaming mode offers superior spatially resolved data and is more suited for mapping neurophysiological features within deep structures (van Wijk et al., 2022) and integrating connectomic data (Oswal et al., 2021). Therefore, we anticipate the identification or development of a simple, robust synchronisation method for this mode in the future, possibly facilitated by further advancements in Percept PC hardware and software.

### 4.5. Summary

Our results confirm the feasibility of using Percept PC with SenSight leads for combined telemetric streaming and MEG recordings. Advanced offline artefact removal techniques are likely to further improve MEG data quality. We also show that the novel OPM MEG sensors are more resilient to monopolar stimulation artefact than conventional cryogenic sensors which might open new avenues for application of OPMs for studying DBS effects.

## Code and Data sharing

The code used to generate the spectral plots can be found in https://github.com/vlitvak/meg-phantom-perceptPC. The phantom data will be made freely available via zenodo.org or similar upon the acceptance of the paper. Early access can be provided by request to the corresponding author.

## Acknowledgements

MFH is supported by a Non-Clinical Post Doctorial Fellowship from the Guarantors of Brain. The Wellcome Centre for Human Neuroimaging is supported by core funding from Wellcome [203147/Z/16/Z]. We thank Jennifer Halcome for her help with arranging the Percept PC loan to UCL.

## Investigating the interaction between the OPM modulation signal and monopolar DBS stimulation

In the main manuscript we saw there were additional peaks in the spectrum when using monopolar stimulation with the OPM sensors compared the CTF-MEG SQUID system. We mentioned that many of the peaks could be determined by an interaction of the DBS device and the OPM modulation signal at 923 Hz. Here we show how these peaks can be determined by calculating the beat frequencies between nearby spectral peaks to 923 Hz.

**Supplementary Figure S1.**
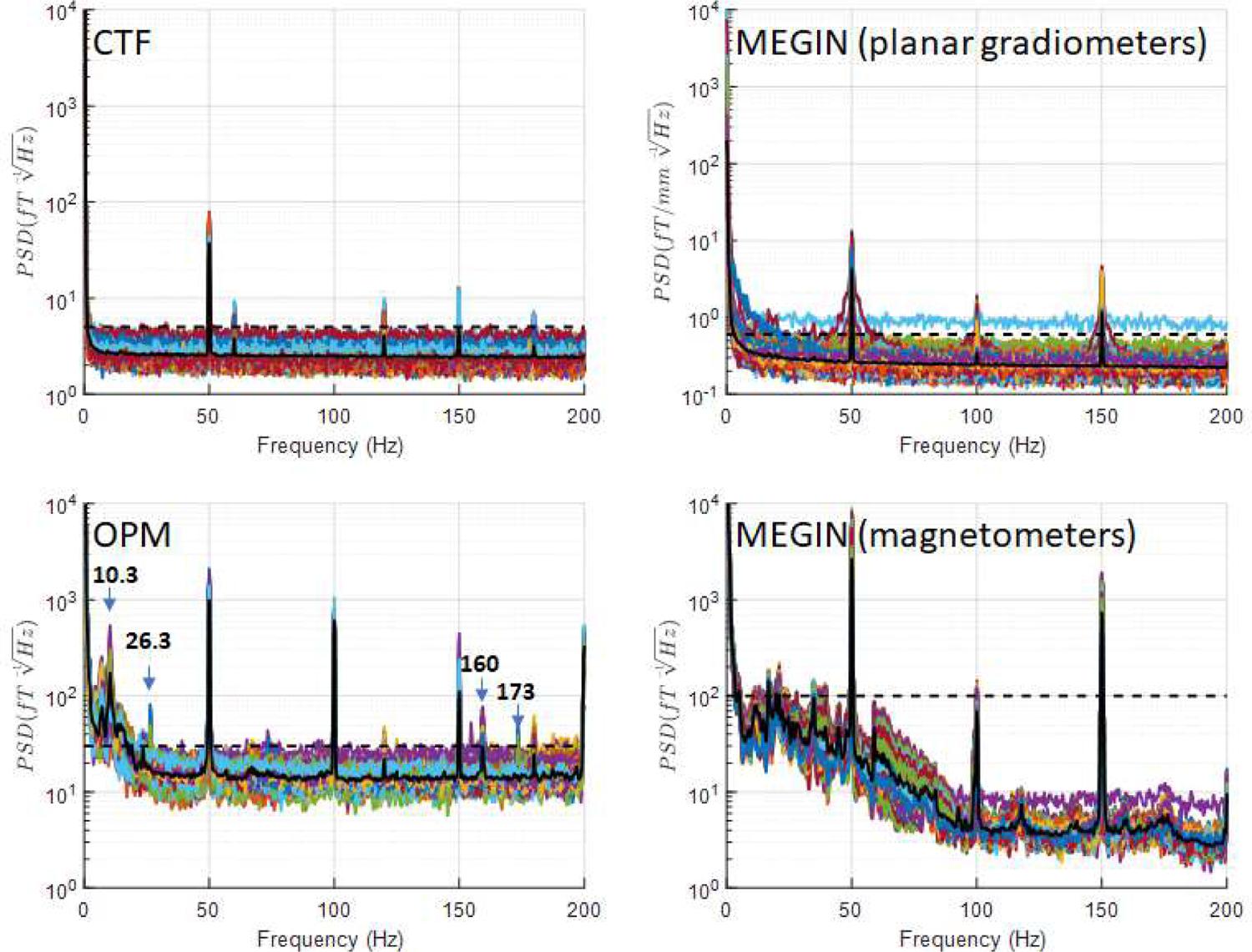
Power spectral density (PSD) for empty room recordings. Recordings were conducted without IPG and experimenter presence. The solid black line shows the average PSD across sensors and the dashed horizontal line shows the upper limit on empty room noise level that is drawn on all the figures for reference. This was 5 fT/√Hz for CTF, 30 fT/√Hz for OPM, 0.6 fT/[mm √Hz] for MEGIN planar gradiometers and 100 fT/√Hz for MEGIN magnetometers. Prominent peaks specific to the OPM system are marked with arrows and the corresponding frequencies are indicated.

Figure S2 shows the spectrum of the OPM recording during monopolar stimulation 180 Hz either side of the modulation frequency (923 Hz; labelled peak 08). 11 high amplitude peaks (and the modulation signal peak) have been identified by the vertical dashed lines. Note the threshold for selection is arbitrary but we selected any peaks above 15 fT/√Hz. Also note the QuSpin OPM sensors have 1^st^ order low-pass filter from the vapour cell in the OPM starting at ∼130 Hz and a second additional hardware-based low-pass elliptical filter at 500 Hz, so these peaks are considerably larger in amplitude than measured here.

**Figure S2:**
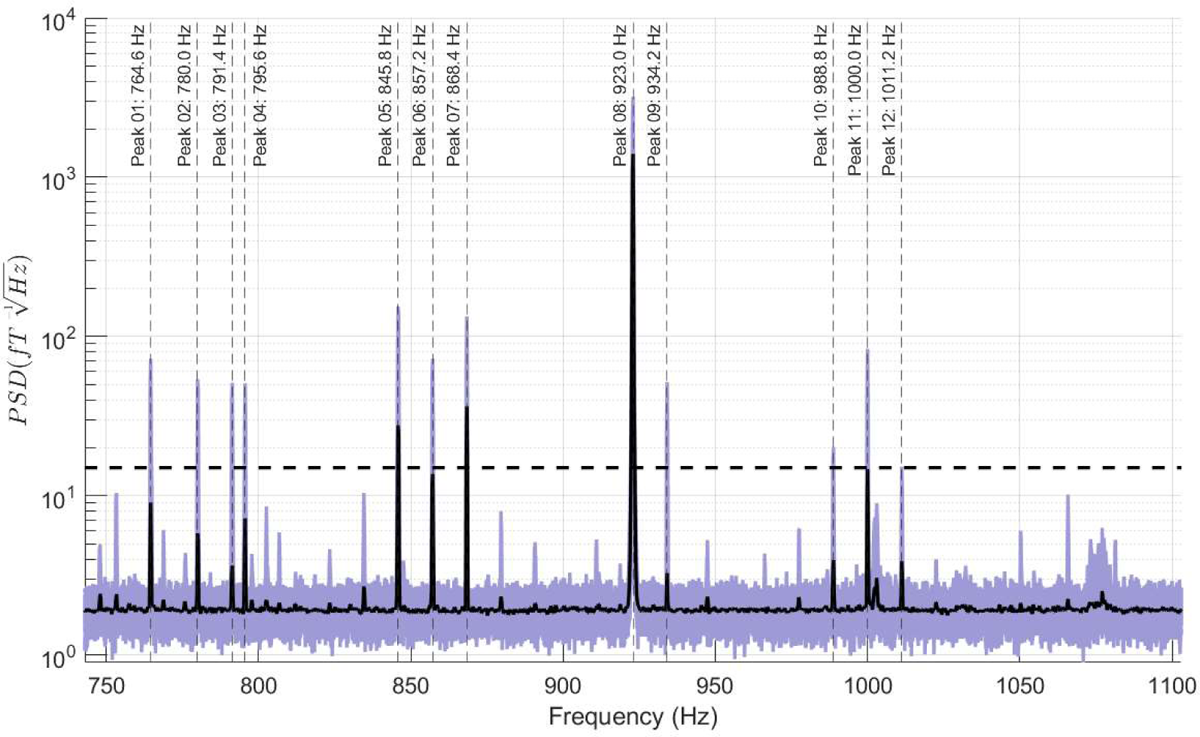
Spectrum of OPM sensor data 180 Hz either side of the modulation frequency (923 Hz) during monopolar stimulation. Here 11 peaks alongside the modulation frequency have been highlighted (vertical dashed lines). Purple lines represent the spectra of individual OPMs, the solid black line is the median spectral density (across all sensors) and the horizontal black dashed line represents 15 fT/√Hz, the threshold used to select peak frequencies for further analyses.

For our first assumption is that the difference in frequency between each of the 11 peaks and the modulation frequency would be represented in the low-frequency spectra. Figure S3 shows the spectrum of the OPM recordings between 0.1—180 Hz. There are multiple clear peaks which were not present in when stimulation is not occurring (see main manuscript). Here we have overlaid the 11 lines representing the expected beat frequencies (vertical dashed lines). We observe that these lines coincide with large peaks in the spectrum supporting that the idea the DBS is beating with the modulation signal of the OPMs. However, this does not explain all the peaks, so we also looked at the difference in frequency between each of the 11 high-frequency peaks to see if these predicted any other features in the spectrum. In Figure S3 we have overlaid beat frequencies which coincided within 0.2 Hz of a peak in the spectrum (the spectral resolution of the fast Fourier transform used). 39 beat frequencies (out of a possible 55) were within 0.2 Hz of a peak, but for clarity we have overlaid 18 (for example two predicted frequencies were at 11.2 Hz and 11.4 Hz)

**Figure S3:**
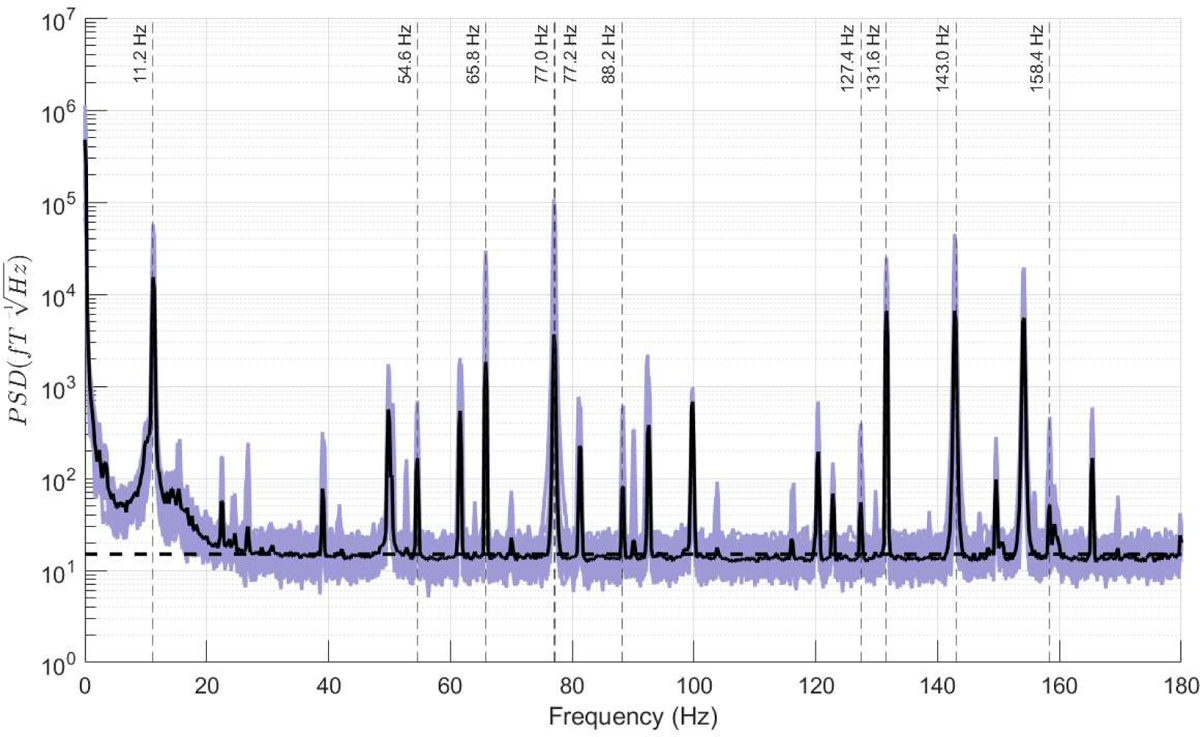
0.1-180 Hz spectrum of the OPM sensors during monopolar stimulation. Overlaid as vertical dashed lines are the frequency differences between the high frequency peaks from Fig. S2 and the modulation frequency. The overlaid frequencies coincide with 11 peaks in the spectrum. Individual sensor spectra are in purple and the median spectra is a solid black line.

**Figure S4:**
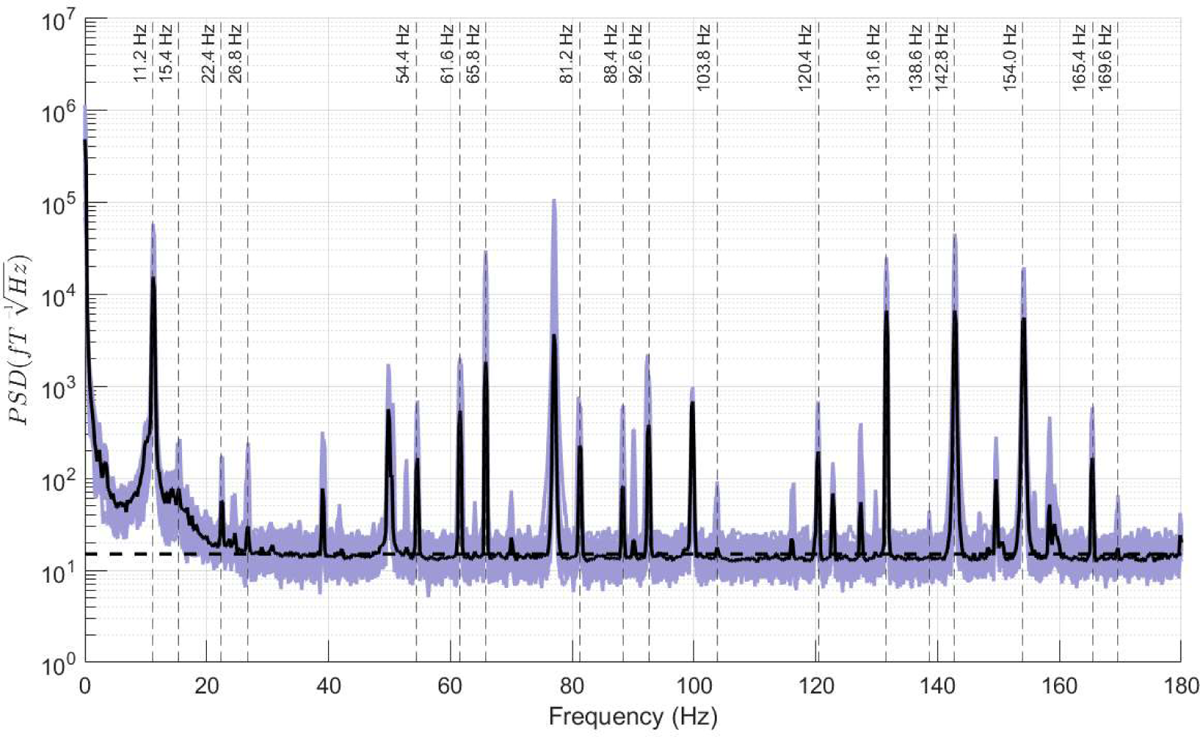
0.1-180 Hz spectrum of the OPM sensors during monopolar stimulation. Overlaid as vertical dashed lines are the frequency differences between the high frequency peaks from Fig. S2, which coincide with large peaks in the low frequency spectrum. Individual sensor spectra are in purple and the median spectra is a solid black line.

